# NVX-CoV2373 vaccination induces functional SARS-CoV-2–specific CD4^+^ and CD8^+^ T cell responses

**DOI:** 10.1101/2022.04.08.487674

**Authors:** Carolyn Rydyznski Moderbacher, Christina Kim, Jose Mateus, Joyce Plested, Mingzhu Zhu, Shane Cloney-Clark, Daniela Weiskopf, Alessandro Sette, Louis Fries, Gregory Glenn, Shane Crotty

## Abstract

NVX-CoV2373 is an adjuvanted recombinant full-length SARS-CoV-2 spike trimer protein vaccine demonstrated to be protective against COVID-19 in efficacy trials. Here we demonstrate that vaccinated subjects made CD4^+^ T cell responses after one and two doses of NVX-CoV2373, and a subset of individuals made CD8^+^ T cell responses. Characterization of the vaccine-elicited CD8^+^T cells demonstrated IFN*γ* production. Characterization of the vaccine-elicited CD4^+^ T cells revealed both circulating T follicular helper cells (cT_FH_) and T_H_1 cells (IFN*γ*, TNFα, and IL-2) were detectable within 7 days of the primary immunization. Spike-specific CD4^+^ T cells were correlated with the magnitude of the later SARS-CoV-2 neutralizing antibody titers, indicating that robust generation of CD4^+^ T cells, capable of supporting humoral immune responses, may be a key characteristic of NVX-CoV2373 which utilizes Matrix-M™ adjuvant.

## INTRODUCTION

The ongoing COVID-19 pandemic caused by severe acute respiratory syndrome coronavirus 2 (SARS-CoV-2) has caused immense morbidity and mortality. This global crisis has been met with an unprecedented surge in vaccine development and implementation. The various COVID-19 vaccines that have been approved for use in humans have vastly improved health outcomes to COVID-19 disease, but the important hurdle of providing COVID-19 vaccines to every person in the world remains.

NVX-CoV2373 by Novavax is composed of recombinant full-length, stabilized prefusion spike protein homotrimers which form approximately 30nm nanoparticles based on hydrophobic interaction with a central polysorbate-80 micelle (1). The vaccine antigen, based on the ancestral Wuhan-Hu-1 strain of SARS-CoV-2, is formulated with Matrix-M™ adjuvant (2). The vaccine is stable at 2-8 °C, making it amenable for deployment to regions where refrigeration is limited, increasing vaccine access for populations where other COVID-19 vaccines are not easily distributed or stored. NVX-CoV2373 has demonstrated efficacy in phase 2b and 3 clinical trials in the UK (3), South Africa (4), and a large phase 3 trial in the United States and Mexico (5). NVX-CoV2373 has emergency use authorization in multiple countries around the globe (6). A more detailed understanding of the immune response to this vaccine is important for the ongoing global efforts to combat COVID-19 infections and deaths, especially in the wake of new and emerging variants (7). Here, we set out to determine the T cell response to the NVX-CoV2373 vaccine, utilizing samples from individuals enrolled in a phase 1/2a clinical trial who received 5 μg NVX-CoV2373 protein adjuvanted with Matrix-M™ (1, 8).

The presence of T cells capable of recognizing SARS-CoV-2 epitopes in SARS-CoV-2 uninfected individuals was first reported in May of 2020 (9–13). These cross-reactive T cells were later confirmed to be memory T cells, some of which were generated in response to infections with commonly circulating human coronaviruses (HCoVs OC43, 229E, NL63, and HKU1) and capable of cross-reacting with shared SARS-CoV-2 epitopes (10, 11, 14). This finding led to speculation that these cells could have an impact on an individual’s immune response to SARS-CoV-2 infection. Theories proposed that these pre-existing T cells could be beneficial or detrimental for immunity to SARS-CoV-2 (15). However, at the time it was unknown what biological effect these cross-reactive memory CD4^+^ T cells might have in the context of COVID-19 vaccination. Therefore, in addition to measuring human CD4^+^ and CD8^+^ T cell responses to NVX-CoV2373, we explored the effect of cross-reactive CD4^+^ T cells on the human immune response to the NVX-CoV2373 vaccine.

## RESULTS

### NVX-CoV2373 induces SARS-CoV-2–specific CD4^+^ T cells

Peripheral blood mononuclear cells from 27 volunteers immunized with 5 μg of NVX-CoV2373 on days 0 and 21, 5 volunteers immunized with 5ug of NVX-CoV2373 on day 0 and placebo on day 21, and 4 recipients of placebo, were isolated from blood samples taken at days 0, 7, and 28. The 5 volunteers that received the placebo dose at day 21 were only included for analysis at day 0 and 7 in all analyses reported in this study. SARS-CoV-2 spike-specific CD4^+^ T cells were measured by activation induced marker (AIM) assay (surface CD40L^+^ (sCD40L) OX40^+^, **Fig 1A** and **Fig S1A**). At day 0, 16% of donors (5/32) had detectable SARS-CoV-2 spike-specific CD4^+^ T cells, detectable above the LOQ for the assay, indicative of pre-existing T cell memory (**Fig 1B,C**). Notably, by day 7 after the 1^st^ immunization 50% of donors (16/32) had developed spike-specific CD4^+^ T cell responses (**Fig 1B,C**). A majority of donors (81%, 22/27) exhibited high levels of SARS-CoV-2 spike-specific CD4^+^ T cells 7 days post-2^nd^ immunization (28 days post-1^st^ immunization, **Fig 1B,C**). There was no significant difference in the magnitude of the antigen-specific CD4^+^ T cell response comparing 7 days after the 1^st^ or 2^nd^ immunization (**Fig 1B,C**). However, the proportion of individuals mounting a detectable SARS-CoV-2 spike response post-2^nd^ immunization was significantly increased relative to post-1^st^ immunization (Fisher’s exact test, p=0.015). CD4^+^ T cell response results were comparable when using OX40^+^4-1BB^+^ AIM (**Fig S1B**), or when calculated by stimulation index (**Fig S1E**). To verify the spike T cell response was vaccine-specific, a peptide megapool (MP) containing predicted SARS-CoV-2 class II epitopes spanning the entire SARS-CoV-2 proteome sans spike, (hereafter referred to as “non-spike”), as well as a CMV MP containing class I and II epitopes, were run in parallel for a subset of donors. There were no significant differences in the CD4^+^ T cell response to non-spike or CMV MPs across the study timepoints, validating the NVX-CoV2373 vaccine spike-specific CD4^+^ T cell response detected (**Fig S1C-D**).

**Figure 1.**
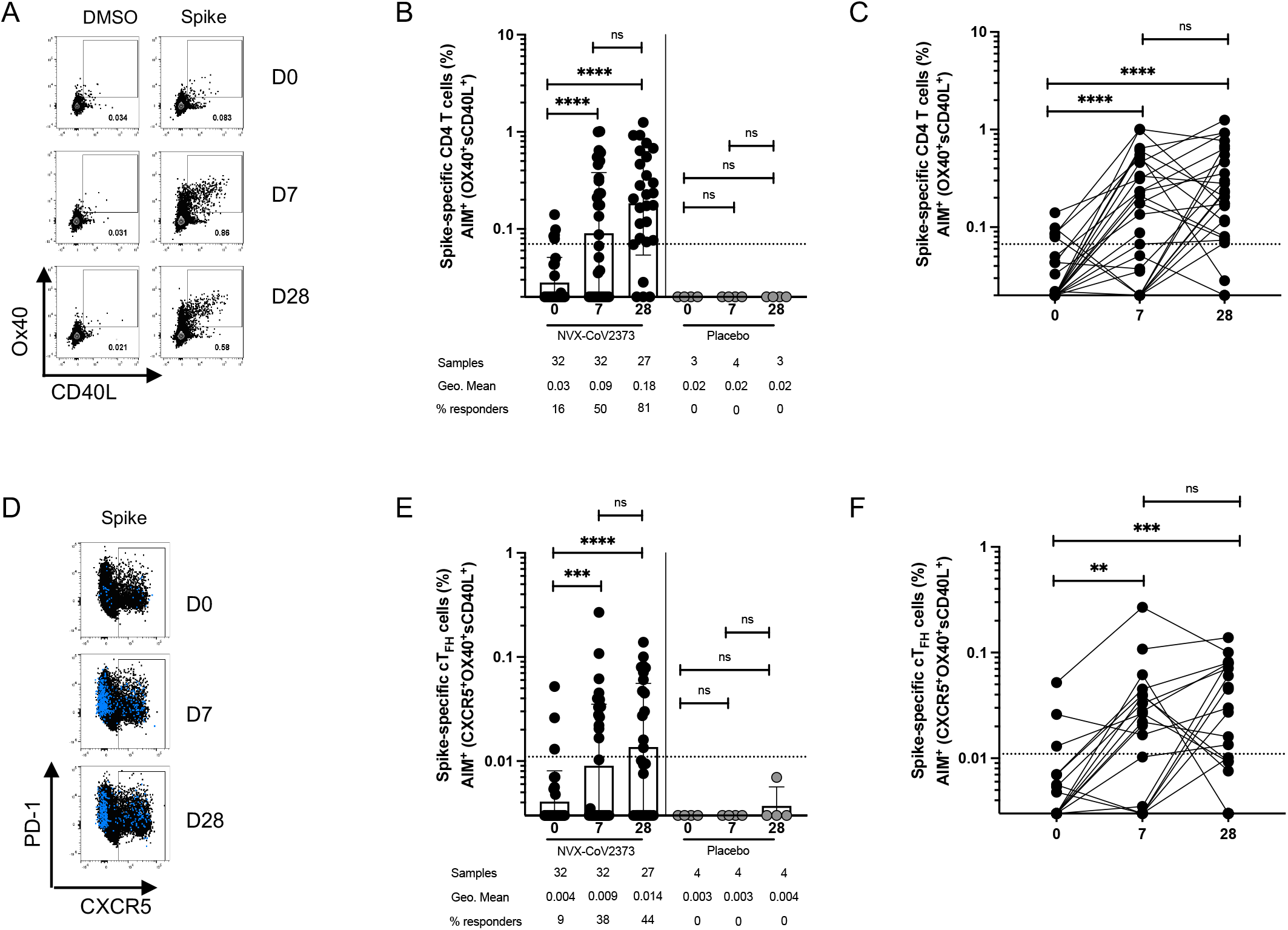
Spike-specific CD4^+^ T cells following NVX-CoV2373 vaccination. **(A)** Representative FACS plots of AIM^+^ (sCD40L^+^OX40^+^) CD4^+^ T cells at D0, D7 and D28 post-vaccination. **(B-C)** Spike-specific AIM^+^ CD4^+^ T cell responses in vaccinees (black dots) and placebo controls (grey dots). The same data is graphed as grouped **(B)** and paired comparisons **(C). (D)** Representative FACS plots of AIM^+^ CD4^+^ CXCR5^+^ cT_FH_ cells (blue dots are AIM^+^ CD4^+^ T cells overlaid on total CD4^+^ T cells in black). **(E-F)** Spike-specific AIM^+^ cT_FH_ cell responses in vaccinees and placebo controls. The same data is graphed as grouped **(E)** and paired comparisons **(F)**. Dotted line indicates limit of quantitation (LOQ) for the assay, and is calculated as the geometric mean of all sample DMSO wells multiplied by the geometric SD factor. % responders are calculated as responses ≥ LOQ divided by the total samples in the group.

Functionalities of NVX-CoV2373-induced CD4^+^ T cell responses were assessed by identifying spike-specific circulating follicular helper T cells (cT_FH_) and by intracellular cytokine staining (ICS) of spike-specific CD4^+^ T cells. T_FH_ cells are crucial for antibody responses following infection or vaccination. Similar to total AIM^+^ CD4^+^ T cells, AIM^+^ cT_FH_ cells (CXCR5^+^ AIM^+^ (sCD40L^+^OX40^+^) CD4^+^ T cells) were significantly increased relative to day 0 in 36% of vaccinees (12/32) one-week post-1^st^ immunization and 44% (12/27) 7 days post-2^nd^ immunization (**Fig 1D-F**). There was no significant difference in the magnitude of spike-specific cT_FH_ cells after the 1^st^ compared to the 2^nd^ immunization, or the proportion of individuals that developed spike-specific cT_FH_ cells (**Fig 1E-F**, and Fisher’s exact test, p=0.61).

Cytokine production by NVX-CoV2373-induced spike-specific CD4^+^ T cells was assessed via ICS. Significant increases in spike-specific IFNγ^+,^, TNFα^+^ and IL-2^+^ CD4 T cells were observed 7 days post-1^st^ immunization (**Fig 2A-C**). IFNγ^+^, TNFα^+^ and IL-2^+^ secreting CD4 T cell frequencies were further increased after the 2^nd^ immunization (**Fig 2A-C**). IL-17α^+^, IL-4^+^, or IL-10^+^ spike-specific cells were not detected (**Fig S2A-C**). Spike-specific IFNγ^+^ intracellular CD40L^+^ (iCD40L) double-positive CD4^+^ T cells were significantly increased after both 1^st^ and 2^nd^ immunizations relative to baseline (**Fig 2D-E**), consistent with the IFNγ^+^ single gating (**Fig 2A**), and included the vast majority of the IFNγ^+^-positive cells (**Fig S2A**). The total cytokine^+^ CD4^+^ T cell response (sum of iCD40L^+^ cells expressing Granzyme B (GzmB), IFN_γ′_, TNFα or IL-2), was significantly increased post-1^st^ and post-2^nd^ relative to baseline (**Fig 2F**). The overall cytokine profile was indicative of a T_H_1 response, appropriate for antiviral immunity. Polyfunctional CD4^+^ T cell cytokine responses were also induced by NVX-CoV2373 vaccination. The proportion of spike-specific CD4^+^ T cells exhibiting 2, 3, 4 or 5 functions (iCD40L, GzmB, IFNγ, TNFα, IL-2) was increased post-1^st^ and post-2^nd^ immunization relative to baseline (**Fig 2G-I**), and a larger proportion of spike-specific CD4^+^ T cells exhibited 3-5 functions one week after the 2^nd^ immunization (35%) relative to spike-specific CD4^+^ T cells after the 1^st^ immunization (19%) (**Fig 2J**). SARS-CoV-2 non-spike IFNγ^+^iCD40L^+^ CD4^+^ T cell frequencies remained unchanged after immunization, as expected (**Fig S2E**). Together, these data suggest that the NVX-CoV-2373 vaccine induces multifunctional spike-specific CD4^+^ T cells composed of classical T_H_1 T cells and cT_FH_ cells necessary for supporting antiviral and antibody responses.

**Figure 2.**
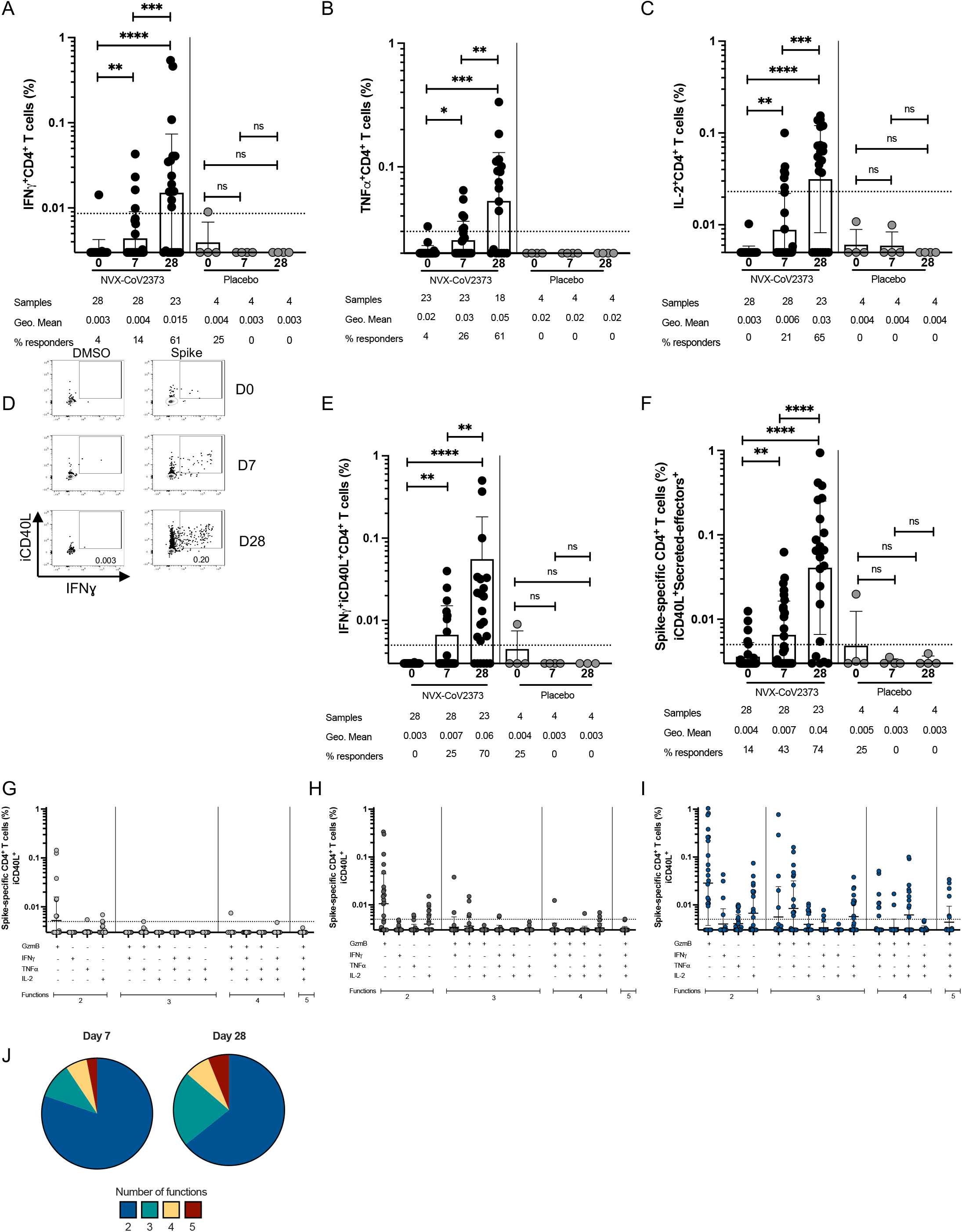
Cytokine-producing spike-specific CD4^+^ T cell responses following NVX-CoV2373 vaccination. Proportion of **(A)** IFNƔ^+^, **(B)** TNFα^+^ and **(C)** IL-2^+^ spike-specific CD4^+^ T cells detected following peptide stimulation. **(D)** Representative FACS plots and **(E)** proportion of IFNƔ^+^ intracellular CD40L^+^ (iCD40L^+^) responses in spike-specific CD4^+^ T cells at days 0,7, and 28 post-vaccination. **(F)** Proportion of spike-specific CD4^+^ T cells expressing iCD40L and producing IFNƔ^+^, TNFα^+^, IL-2^+^, or GzmB (“secreted-effector^+^”). Predominant multifunctional profiles of spike-specific CD4^+^ T cells with one, two, three, four or five functions were analyzed at **(G)** D0, (**H**) D7, and **(I)** D28 post vaccination. **(J)** Pie charts depicting the proportion of spike-specific CD4^+^ T cells exhibiting 2, 3, 4, or 5 functions at day 7 and day 28 post-immunization. Dotted line indicates LOQ for the assay, and is calculated as the geometric mean of all sample DMSO wells multiplied by the geometric SD factor. % responders are calculated as responses ≥ LOQ divided by the total samples in the group.

### NVX-CoV2373 induces SARS-CoV-2–specific CD8^+^ T cells

Spike-specific CD8^+^ T cell responses following 1^st^ and 2^nd^ immunizations with the NVX-CoV2373 vaccine were tested by AIM and ICS assays (**Fig 3A and S3A, E**). No donors had detectable CD8^+^ T cells by AIM at baseline (CD69^+^4-1BB^+^, **Fig 3A-C**). Seven days post-1^st^ immunization, 9% of donors (3/32 donors) had a modest spike-specific CD8^+^ T cell response detected by AIM. One-week post-2^nd^ immunization, 20% of donors had developed a spike-specific CD8^+^ T cell response (5/27 donors, **Fig 3A-C**). Stimulation indices similarly showed significant NVX-CoV2373 vaccine-induced CD8^+^ T cell AIM^+^ responses post-1^st^ and post-2^nd^ immunization (**Fig S3D**). There was no significant difference in the amplitude of CD8^+^ T cell response between post-1^st^ and post-2^nd^ immunization timepoints (**Fig 3C**, and Fisher’s exact test, p=0.5), but the majority of donors who had developed CD8^+^ T cell responses 7 days post-1^st^ immunization retained these responses post-2^nd^ immunization, and were joined by an additional cohort of CD8^+^ responders after the second dose. There was no difference in the CD8^+^ T cell frequency to SARS-CoV-2 non-spike or CMV MPs across all timepoints assessed, as expected (**Fig S3B**,**C**).

**Figure 3.**
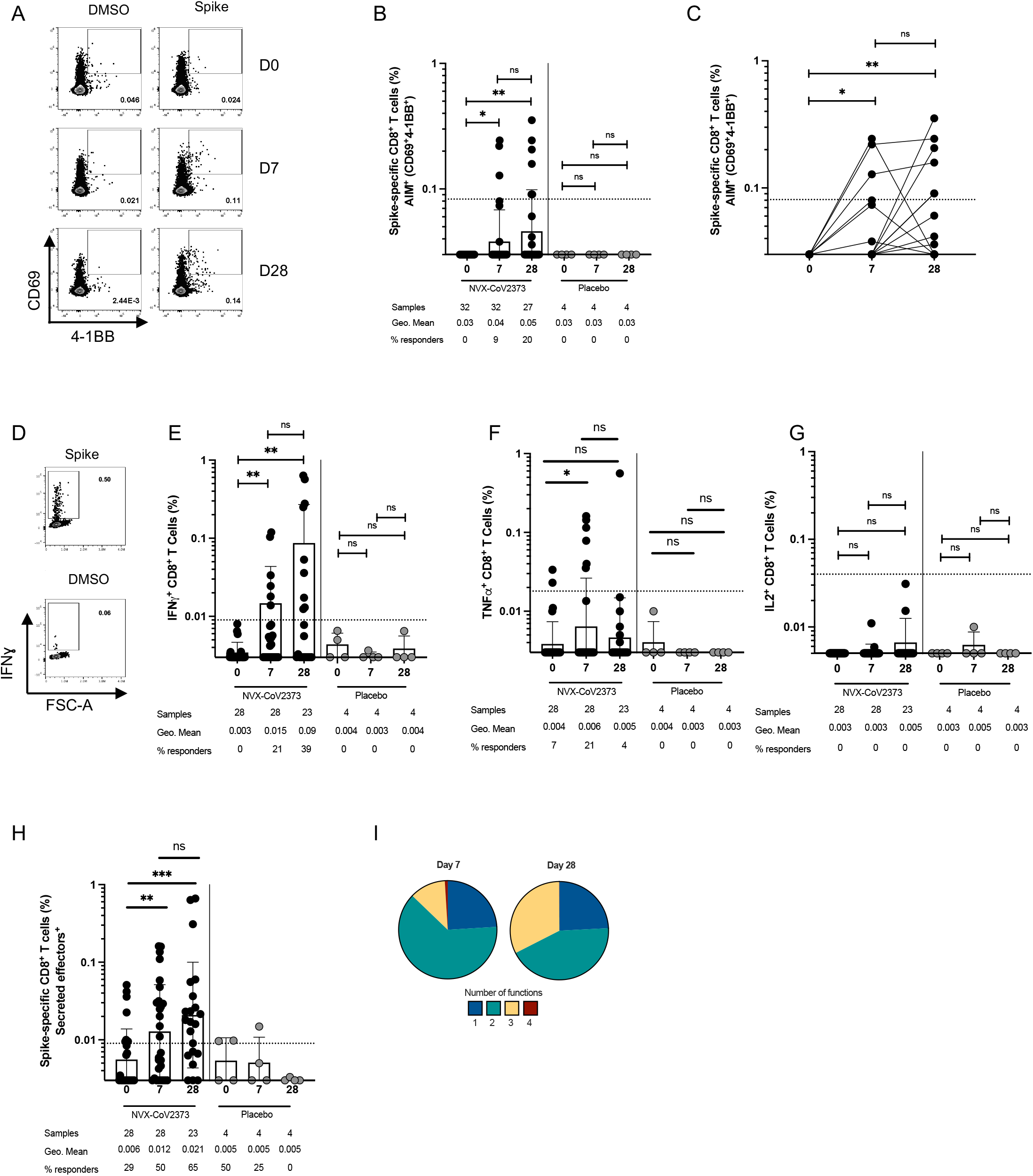
Spike-specific CD8^+^ T cells are induced following NVX-CoV2373 vaccination. **(A)** Representative FACS plots of AIM^+^ (CD69^+^4-1BB^+^) CD8^+^ T cells at D0, D7 and D28 post-vaccination. **(B-C)** Spike-specific AIM^+^ CD8^+^ T cells responses in vaccinees (black dots), placebo controls (grey dots) and convalescent COVID-19 donors (blue dots). **(D)** Representative FACS plots of IFNƔ responses in total spike-specific CD8^+^ T cells. IFNƔ^+^ **(E)**, TNFα^+^ **(F)** and IL-2^+^ **(G)** Spike-specific CD8^+^ T cell responses in vaccinees, placebo controls and convalescent COVID-19 controls. **(H)** Secreted-effector^+^ spike-specific CD8^+^ T cells (sum of CD8^+^ T cells expressing any combination of IFN, TNF, or IL-2 or GzmB, excluding GzmB single positives). **(I)** Pie charts depicting the proportion of spike-specific CD8^+^ T cells exhibiting 1, 2, 3, or 4 functions at day 7 and day 28 post-immunization (combinations of IFN, TNF, IL-2 or GzmB, excluding GzmB single positives). Dotted line indicates limit of sensitivity for the assay, and is calculated as the geometric mean of all sample DMSO wells multiplied by the geometric SD factor. % responders are calculated as responses ≥ LOQ divided by the total samples in the group.

SARS-CoV-2 spike-specific CD8^+^ T cell functionality was assessed via ICS. Spike-specific IFNγ^+^ CD8^+^ T cells were detected in a small subset of donors post-1^st^ immunization (11%, 3/28 donors) and increased after the 2^nd^ immunization (26%, 6/23 donors. **Fig 3D,E**). Similar increases in IFNγ^+^ CD8^+^ T cells after 1^st^ and 2^nd^ immunizations were also observed when the data was plotted as stimulation index. (**Fig S3F**). TNFα was modestly increased after the 1^st^ (21% 6/28 donors), but not 2^nd^ immunization (**Fig 3F**). IL-2^+^ single-positive CD8^+^ T cells were not detected (**Fig 3G**). The total cytokine^+^ CD8^+^ T cell response (sum of CD8^+^ T cells expressing any combination of IFNγ, TNFα, IL-2 or GzmB, excluding GzmB single positives), was significantly increased after both 1^st^ and 2^nd^ immunizations relative to baseline (**Fig 3H**). Polyfunctional CD8^+^ T cell cytokine responses were also induced by NVX-CoV2373 vaccination. The proportion of spike-specific CD8^+^ T cells exhibiting 2, 3 or 4 functions (GzmB, IFNγ, TNFα, IL-2) was increased post-1^st^ and post-2^nd^ immunization relative to baseline (**Fig S3G-I**). The proportion of spike-specific CD8+ T cells positive for 3 functions was increased after the 2^nd^ immunization (29%) relative to the 1^st^ immunization (9%) (**Fig 3I**). In sum, spike-specific CD8^+^ T cell responses were induced in some subjects following NVX-CoV2373 vaccination, as measured by two experimental approaches.

### Relationship between T cell and antibody responses following NVX-CoV2373 vaccination

Samples were analyzed for spike IgG titers via ELISA and SARS-CoV-2 neutralization activity using both microneutralization and hACE2 binding inhibition assays (**Fig S4A-C**). Spike IgG and neutralizing antibodies were induced following the 1^st^ immunization and further enhanced upon the 2^nd^ immunization (**Fig S4A-C**). Spike-specific CD4^+^ T cells and antibody responses were assessed for relationships. Spike-specific (AIM^+^) CD4^+^ T cells did not significantly correlate with spike IgG titers following the 1^st^ immunization, but were significantly associated following the 2^nd^ immunization (**Fig 4D and S4D**). However, AIM^+^ CD4^+^ T cells were significantly associated with neutralizing antibodies after both 1^st^ and post-2^nd^ immunization (**Fig 4A, B,C**). Spike-specific cT_FH_ cell frequencies did not measurably correlate with spike IgG or neutralizing antibodies (**Fig S4E-H**). AIM^+^ CD4^+^ T cell and AIM^+^ CD8^+^ T cell responses were significantly correlated at both 7 days post-1^st^ and 7 days post-2^nd^ immunization (**Fig 4D,E**). Together, these data demonstrate the ability of the NVX-CoV2373 vaccine to elicit antibody, CD4^+^ T cell and CD8^+^ T cell responses against SARS-CoV-2 infection.

**Figure 4.**
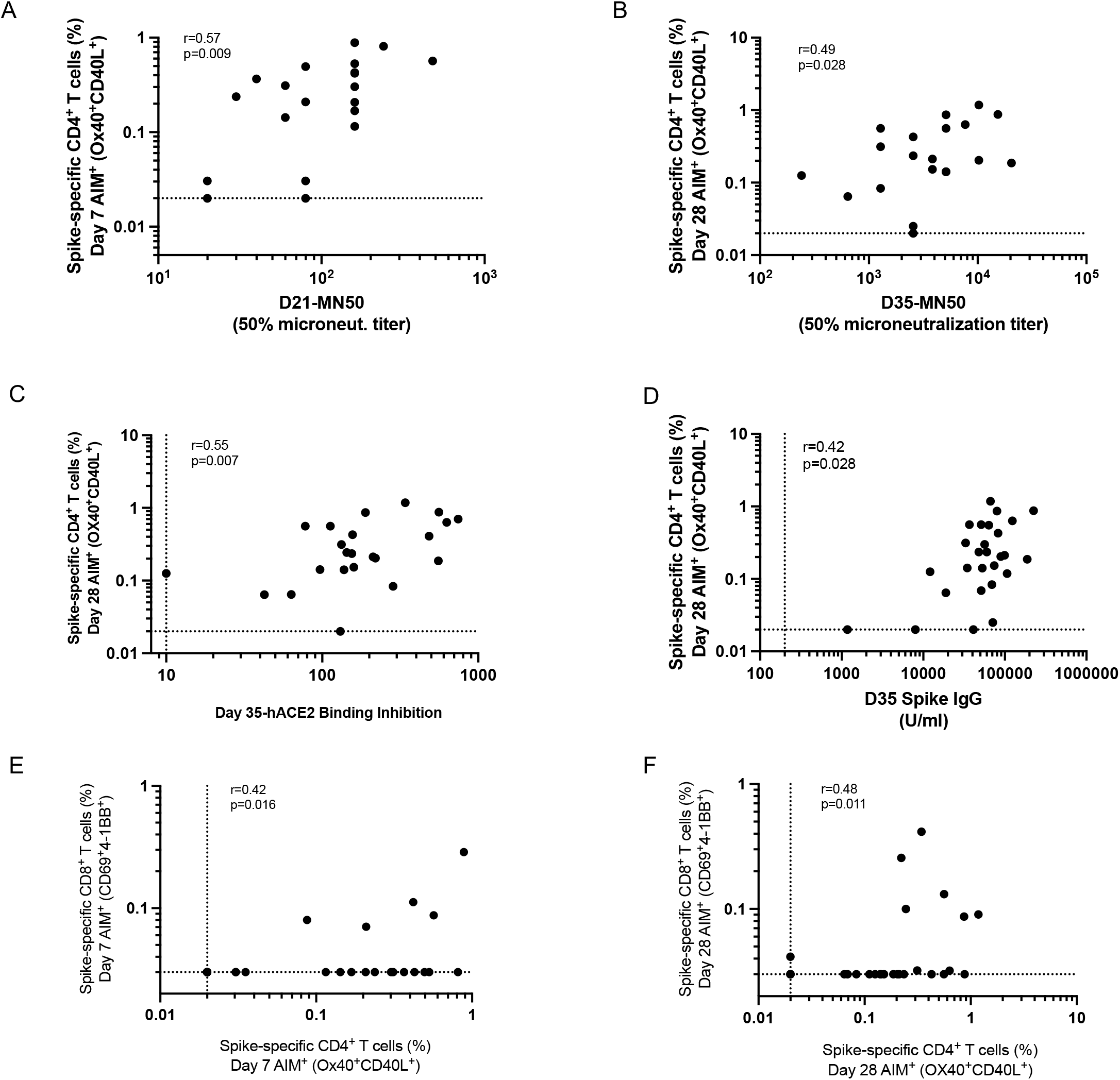
Association between T cell and antibody responses following NVX-CoV2373 vaccination. Correlation between D7 SARS-CoV-2 spike-specific AIM^+^ CD4^+^ T cells and **(A)** day 21 microneutralization titers or **(B)** D28 SARS-CoV-2 spike-specific AIM^+^ CD4^+^ T cells and D35 neutralization titers. Correlation between D28 SARS-CoV-2 Spike-specific AIM^+^ CD4^+^ T cells and **(C)** day 35 hACE2 binding inhibition or **(D)** D35 Spike IgG. Correlation between spike specific AIM^+^ CD4^+^ cells and AIM^+^ CD8^+^ cells at **(E)** D7 and **(F)** D28.

### Pre-existing T cell immunity did not impact NVX-CoV2373 vaccine responses

Within the cohort, 10 donors had detectable SARS-CoV-2 spike-specific CD4^+^ T cell responses at baseline (**Fig 5A**). These donors self-reported no previous SARS-CoV-2 infection or vaccination and were recruited in a time frame when local SARS-CoV-2 seroprevalence was very low. Therefore, these were likely cross-reactive memory T cells, induced via infection with endemic coronaviruses or other infections. Whether such pre-existing cross-reactive cells contribute to, detract from, or have no influence on COVID-19 vaccine responses was undetermined. Thus, we explored this topic in the cohort of NVX-CoV2373 vaccinees. Donors were separated into two groups based on the presence or absence of cross-reactive T cells at baseline (AIM^+^ CD4^+^ T cell response above the LOD for the assay [0.02%]). There was no significant difference in the magnitude or frequency of the spike-specific (AIM^+^) CD4^+^ T cell response post-1^st^ or post-2^nd^ immunization between donors with and without pre-existing crossreactive memory CD4^+^ T cells (**Fig 5A**). Similarly, we observed no significant differences between the magnitude of AIM^+^ cT_FH_ CD4^+^ T cells or AIM^+^ CD8^+^ T cells based upon pre-existing crossreactive memory CD4^+^ T cells (**Fig 5B,C**). Subjects with cross-reactive AIM^+^ CD4^+^ T cells also did not exhibit any significant differences in the magnitude of spike-specific cytokine^+^ CD4^+^ or CD8^+^ T cells (**Fig 5D,E**). These results were similar when we employed a more rigorous threshold for pre-existing T cell immunity, assessing samples with an AIM^+^ CD4^+^ T cell response above the LOQ for the assay (0.067%) (**Fig S5A-E**). Therefore, individuals with crossreactive memory CD4^+^ T cells do not appear to be more or less likely to make a stronger immune CD4^+^ T cell response following two doses of the NVX-CoV2373 vaccine, although this conclusion is based on limited data.

**Figure 5.**
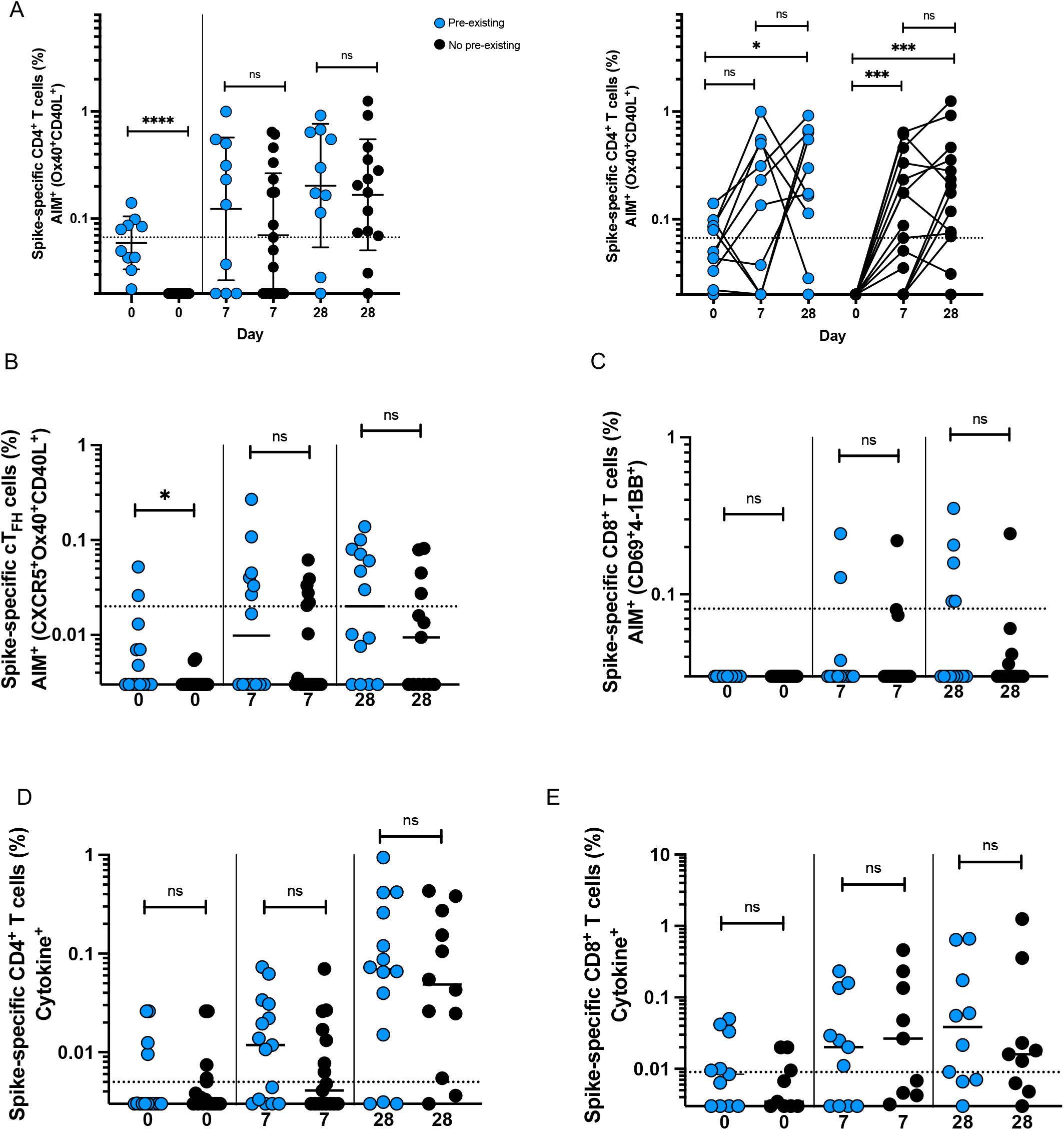
Pre-existing T cell immunity does not impact NVX-CoV2373 vaccine responses. **A)** Spike-specific AIM^+^ CD4^+^ T cells at baseline and post-vaccination grouped according to donors with detectable AIM^+^ CD4^+^ T cell responses at D0 (blue dots) or no detectable AIM^+^ CD4^+^ T cell responses at D0 (black dots). Effect of pre-existing AIM^+^ CD4^+^ T cells on **(B)** spike specific AIM^+^ cT_FH_ cells, **(C)** spike-specific AIM^+^ CD8^+^ T cells, **(D)** total Spike-specific CD4^+^ T cell and **(E)** spike-specific CD8^+^ T cell cytokine production. Dotted line indicates limit of sensitivity for the assay, and is calculated as the geometric mean of all sample DMSO wells multiplied by the geometric SD factor.

## DISCUSSION

Understanding COVID-19 vaccine-induced immunity is crucial for implementing immunization regimens that can be used to mitigate the ongoing SARS-CoV-2 pandemic in various regions of the world and various populations. Samples from 32 healthy adults were analyzed to determine cellular immune responses to the protein-based COVID-19 vaccine, NVX-CoV2373. Immunization with NVX-CoV2373 induced a SARS-CoV-2 spike-specific CD4^+^ T cell response as early as 7 days after the 1^st^ immunization, characterized by a substantial cT_FH_ population, as well as polyfunctional cytokine producing cells dominated by IFNγ, TNFα, and IL-2. Notably, NVX-CoV2373 immunization also promoted modest CD8^+^ T cell responses in a subset of donors. CD4^+^ and CD8 T^+^ cell responses correlated with each other after both 1^st^ and 2^nd^ immunizations, and total AIM^+^ CD4^+^ T cells correlated with neutralizing antibodies.

A combination of AIM assays and ICS were used to characterize SARS-CoV-2 spike-specific CD4^+^ and CD8^+^ T cells induced by NVX-CoV2373 vaccination. Spike-specific CD4^+^ T cells, as measured by AIM assay, could be detected as early as 7 days post-1^st^ immunization. This rapid induction of robust spike-specific CD4^+^ T cell responses could be a result of increased immunogenicity from the Matrix-M™ adjuvant in the NVX-CoV2373 vaccine. Similar to the total spike-specific CD4^+^ T cell response, NVX-CoV2373 also induced a strong cT_FH_ cell response 7 days post-1^st^ immunization. A rapid and sustained cT_FH_ response likely supports a more productive antibody response. We did not observe an increase in the magnitude of the response between the post-1^st^ immunization and post-2^nd^ immunization time points. This appears consistent with findings from multiple groups characterizing responses to two doses of the COVID-19 mRNA vaccines in a similar window of time post-immunization (16, 17).

Functional cytokine profiles of spike-specific CD4^+^ T cells revealed Th1 cytokines produce by the CD4^+^ T cell response induced by NVX-CoV2373 vaccination, including IFNγ, TNFα and IL-2, with minimal IL-10 or Th2 or Th17 cytokine profiles observed. Overall, the cytokine data indicated the vaccine induces a Th1 CD4 T cell response, consistent with earlier reports (1), and mirroring a Th1 bias in the presence of Matrix-M™ that has been reported in several animal models (2, 18). The cytokine responses increased post-2^nd^ immunization, with an increase in polyfunctionality continuing after the 2^nd^ dose of vaccine, indicating further polarization of the CD4^+^ T cell response.

In contrast to live-attenuated or inactivated vaccines, protein-based vaccines have historically proven to be poor inducers of CD8^+^ T cell responses in humans (19–21). However, the data presented here indicate that the NVX-CoV-2373 vaccine does induce modest spike-specific CD8^+^ T cell responses in a subset of individuals. Spike-specific CD8^+^ T cells were identified by AIM. ICS assays showed similar results, with IFNγ^+^ CD8^+^ T cells detected after both 1^st^ and 2^nd^ immunizations. CD8^+^ T cell responses were less polyfunctional than CD4^+^ T cell responses to NVX-CoV2373 vaccination, but several donors did develop multifunctional CD8^+^ T cell responses following their 1^st^ or 2^nd^ dose of NVX-CoV2373. Spike-specific CD4^+^ T cell responses correlated with CD8^+^ T cell responses, suggesting that the CD8^+^ T cell response observed may depend on the CD4^+^ T cell response. Additional studies with a larger cohort of vaccinees including seropositive and seronegative subjects are warranted to validate these findings.

Matrix-M™ adjuvant may be more potent than most adjuvants at inducing CD8^+^ T cell priming, or possibly recalling cross-reactive memory T cells. Amphiphilic saponins, as present in Matrix-M™, are reported to destabilize the endosomal/lysosomal membrane and facilitate entry of antigens into the cytoplasm, a first step in cytosolic catabolism and subsequent presentation in the context of MHC class I to initiate CD8^+^ T cell responses (22).

Cross-reactive CD4^+^T cells, which antedate SARS-CoV-2 or SARS-CoV-2 vaccine exposure and arise in part from endemic beta- and alpha-coronavirus exposure, are capable of recognizing SARS-CoV-2 epitopes, have been of substantial interest (10–15, 23–26). One theory suggests cross-reactive T cells could amplify the vaccine-response, while others have posited that these cells could largely comprise T cells with relatively low-affinity for SARS-CoV-2 and might inhibit higher affinity T cell clones from participating in the immune response (15). Within this cohort, we identified 10 donors who had spike-specific AIM^+^ CD4^+^ T cells at baseline. We identified no effect of pre-existing cross-reactive memory CD4^+^ T cells at baseline on subsequent SARS-CoV-2 spike-specific T cell responses to vaccination with NVX-CoV2373. This is in contrast to reports examining cross-reactive memory CD4^+^ T cells in the context of mRNA COVID-19 vaccines (17, 26). One explanation for this difference between studies could be differences between protein and mRNA vaccine mechanisms of priming of T cell and B cell responses in vivo. Future studies directly comparing pre-existing T cells across different vaccine platforms will be necessary to elucidate any role for these cells in vaccine responses.

Spike-specific CD4^+^ T cells induced by NVX-CoV2373 vaccination correlated with SARS-CoV-2 neutralizing titers. While T_FH_ cells are highly likely to be the CD4^+^ T cell subset responsible for B cell help and neutralizing antibody development after NVX-CoV2373 immunization, spike-specific cT_FH_ cells did not show a statistically significant correlation with neutralizing titers. This could simply be due to sample size, or the greater difficulty quantifying antigen-specific cT_FH_ cells in blood. Additionally, by directly observing GC-T_FH_ cells in lymph nodes of vaccinated individuals, it was observed that spike-specific T_FH_ cell frequencies in lymph nodes correlated with germinal center responses and neutralizing antibodies, but cT_FH_ cell frequencies in the blood did not [(25)]. Lastly, a markedly enhanced antibody response observed after the first dose of NVX-CoV2373 in baseline seropositives who were previously exposed to pandemic SARS-CoV-2 (Madhi S, in press), consistent with memory CD4^+^ T cells and B cells to spike being recalled by NVX-CoV2373 immunization.

Together, these data support the idea that the NVX-CoV2373 vaccine induces a complex immune response consisting of robust and polyfunctional CD4^+^ T cells producing Th1-type cytokines as well as a rapid cT_FH_ cell response capable of supporting a substantial neutralizing antibody response, as well as a modest CD8^+^ T cell response in a subset of donors. Overall, these data show that NVX-CoV2373 induces a relatively broad humoral and cellular immune response against SARS-CoV-2 in humans, and might demonstrate distinctive long-term behavior relative to mRNA vaccines.

## METHODS

### Human Subjects

Peripheral blood mononuclear cells (PBMC) were obtained from subjects in study 2019nCoV-101, a phase I/II clinical trial of NVX-CoV2373 carried out in male and female adult subjects in Australia and the United States. The protocol and consent document were reviewed and approved by ethical review boards for all sites, and all subjects provided written informed consent. Dosing was carried out under supervision of a Data and Safety Monitoring Board. Descriptions of parts I and II of this trial have been published elsewhere (1, 8). Subjects included healthy adults without evidence of preceding SARS-CoV-2 infection and ages 18 to 59 years in part I, with the addition of a 60- to 84-year-old stratum in part II. Donors of peripheral blood mononuclear cell fractions for the studies reported here were selected randomly from among subjects who had adequate specimens at all three specified dates (baseline, 7 days after dose 1 and 7 days after dose 2) and were treated twice with 5µg SARS-CoV-2 rS antigen plus 50µg Matrix-M™ adjuvant at a 21-day interval, as this was the dose and regimen selected to go forward for further clinical development. PBMC from 5 recipients of placebo were included among the study samples in a blinded fashion.

### Flow Cytometry

#### T cell stimulation

For all flow cytometry assays of stimulated T cells, cryopreserved cells were thawed by diluting them in 10mL pre-warmed complete RPMI containing 5% human AB serum (Gemini Bioproducts) in the presence of benzonase (20 μl/10mL) and spun at 1200 rpm for 7 min. Supernatants were carefully removed by pipetting and cells were resuspended in warm medium, counted and apportioned for assays

#### Activation induced cell marker assay

Assays were conducted as previously described (13, 26, 27). Cells were cultured for 24 h in the presence of SARS-CoV-2-specific MPs [1 μg/mL] in 96-well U bottom plates at 1×10^6^ PBMC per well in complete RPMI containing 5% Human AB Serum (Gemini Bioproducts). Prior to addition of peptide MPs, cells were blocked at 37 C for 15 min with 0.5 μg/mL anti-CD40 mAb (Miltenyi Biotec). A stimulation with an equimolar amount of DMSO was performed as negative control, Staphylococcal enterotoxin B (SEB, 1 μg/mL) and stimulation with a combined CD4^+^ and CD8^+^ T cell cytomegalovirus MP (CMV, 1 μg/mL) were included as positive controls. Supernatants were harvested at 24 h post-stimulation for multiplex detection of cytokines. Antibodies used in the AIM assay are listed in Table S4. AIM^+^ gates were drawn relative to the unstimulated condition for each donor. Stimulation index is defined as the background subtracted response to a MP divided by the average DMSO response for that sample. Poor quality samples were identified as samples with an SEB response less than 50% of the median SEB response for all samples, and were excluded from downstream analyses. Values were set to the LOD for the assay if the stimulation index was <2. Stimulation index is calculated as the background subtracted signal in the test condition divided by the average response detected in the DMSO negative control wells for that sample. Limit of quantitation (LOQ) is calculated as the geometric mean of all sample DMSO wells multiplied by the geometric SD factor. % responders are calculated as responses ≥ LOQ divided by the total samples in the group.

#### Intracellular cytokine staining (ICS) assay

Before the addition of MPs, cells were blocked at 37°C for 15 min with 0.5 μg/ml of anti-CD40 mAb, as previously described (reference). PBMCs were cultured in the presence of SARS-CoV-2 MPs (1 μg/ml) for 24 hours at 37°C. In addition, PBMCs were incubated with an equimolar amount of DMSO as a negative control and also CMV MP (1 μg/ml) as a positive control. After 24 hours, Golgi-Plug and Golgi-Stop were added to the culture for 4 hours, as described above. Cells were then washed and surface-stained for 30 min at 4°C in the dark and fixed with 1% of paraformaldehyde (Sigma-Aldrich, St. Louis, MO). LOQ and % responders are calculated as described above.

#### Anti-S IgG ELISAs

Recombinant SARS-CoV-2 S protein was immobilised onto the surface of the 96-well microtiter plates by direct adsorption at 2°C to 8°C, followed by washing and blocking, Diluted reference standard (2-fold dilution series of 12 dilutions starting 1:1000) and human serum samples (3-fold dilution series of 12 dilutions) in assay buffer were then added in duplicate (100 µL per well) to the S protein-coated wells and specific antibodies are allowed to complex with the coated antigen for 2 hours ± 10 minutes at 24°C ± 2°C. After washing, IgG bound to the rSARS-CoV-2 S protein was detected using a horseradish peroxidase (HRP)-conjugated goat anti-human IgG antibody (Southern Biotech) incubated for 1 hour ± 10 minutes at 24°C ± 2°C. After further washing, a colorimetric signal was generated by addition of 100 µL per well 3, 3′,5,5′-tetramethylbenzidine (TMB) substrate for 10 minutes ± 2 minutes at 24°C ± 2°C. The TMB reaction was stopped with 100 µL per well of TMB Stop solution, and absorbance was measured at 450 nm. Anti-rSARS-CoV-2 S protein IgG antibody level in clinical serum samples was quantitated in ELISA unit, EU/mL, by comparison to a reference standard curve. The results were analysed by SoftMax Pro software using a 4-PL curve fit. Each assay run included control plates comprising of positive and negative controls.

#### hACE2 Binding Inhibition Assay

SARS-CoV-2 (rSARS-CoV-2) S protein was immobilised onto the surface of the 96-well microtiter plates by direct adsorption at 2°C to 8°C, followed by washing and blocking, Serial dilutions of human serum samples, including assay quality controls (QCs), were then added to the spike-coated wells and any molecules that could bind to the S protein, presumptively primarily spike-specific antibodies, were allowed to complex with the immobilized S protein (for 1 hour at 24±2°C) After a plate wash step, a fixed concentration of human ACE2 receptor (hACE2) with a polyhistidine-Tag (His-Tag) (SinoBiological) was added to the plate for incubation (1 hour at 24±2°C) during which the hACE2 bound to the S protein residues with binding sites not obstructed by bound antibody. After washing, the hACE2 receptor bound to the S protein was then detected using a mouse anti-His-Tag horseradish peroxidase conjugate (Southern Biotech) and a colorimetric signal generated by the addition of 3,3′,5,5′-TMB substrate. The amount of bound hACE2 detected was inversely proportional to the amount hACE2 binding inhibitors (antibodies) in human serum; inhibitory activity was reported as 50% inhibitory titers based on a 4-PL curve fit (software).

#### SARS-CoV-2 Microneutralization

Microneutralization assays were performed at University of Maryland in the laboratory of Dr. Matthew Frieman using heat-inactivated (56°C x 30 min) sera. Samples were diluted inn duplicate to a base dilution of 1:5 or 1:10, followed by 11 × 1:2 serial dilutions in Dulbecco’s minimal essential medium (DMEM, Quality Biologicals) supplemented with 10% fetal bovine serum (heat inactivated, Sigma), 1% penicillin/streptomycin)(Gemini Bio-products) and 2mM L-glutamine (Gibco) resulting in 100µL per well. The dilution plates were then transferred to a BSL-3 environment and 100µL of pre-diluted SARS-CoV-2 inoculum was added to result in a multiplicity of infection (MOI) of 0.01 upon transfer to 96-well plates. Virus-only and mock-infection well were included in each assay. After incubation of the mixtures at 37°C and 5% CO_2_ for 1 hour, the mixtures were transferred to 96-well plates with confluent VeroE6 cells. The plates were then further incubated at 37°C and 5% CO_2_for 72 hours, followed by examination for cytopathic effect (CPE). The first dilution to show CPE was was reported as the minimum dilution required to inhibit (neutralize) >99% of the inoculum of SRAS-CoV-2 tested.

## ACKNOWLEDGMENTS

Funding support for the initial manufacture of NVX-CoV2373 and the 2019 nCoV-101 study was provided by the Coalition of Epidemic Preparedness Innovations (CEPI) (https://cepi.net) to Novavax, Inc. The current work was funded by the NIH NIAID under awards AI142742 (Cooperative Centers for Human Immunology, CCHI) (A.S., S.C.), and a supplement to NIH AI142742. This work was additionally supported in part by LJI Institutional Funds and the NIAID under K08 award AI135078 (J.M.D.). The contributions of Drs. Cheryl Keech and Neil Formica in conduct of the clinical trial generating the serum and cellular specimens, and Dr. Matthew Frieman (University of Maryland, USA) in conduct of the microneutralization assays, are gratefully acknowledged.

