## Supplemental Figures for "NVX-CoV2373 vaccination induces functional SARS-CoV-2–specific CD4^+^ and CD8^+^ T cell responses"

#### SUPPLEMENTARY FIGURES

### Figure S1

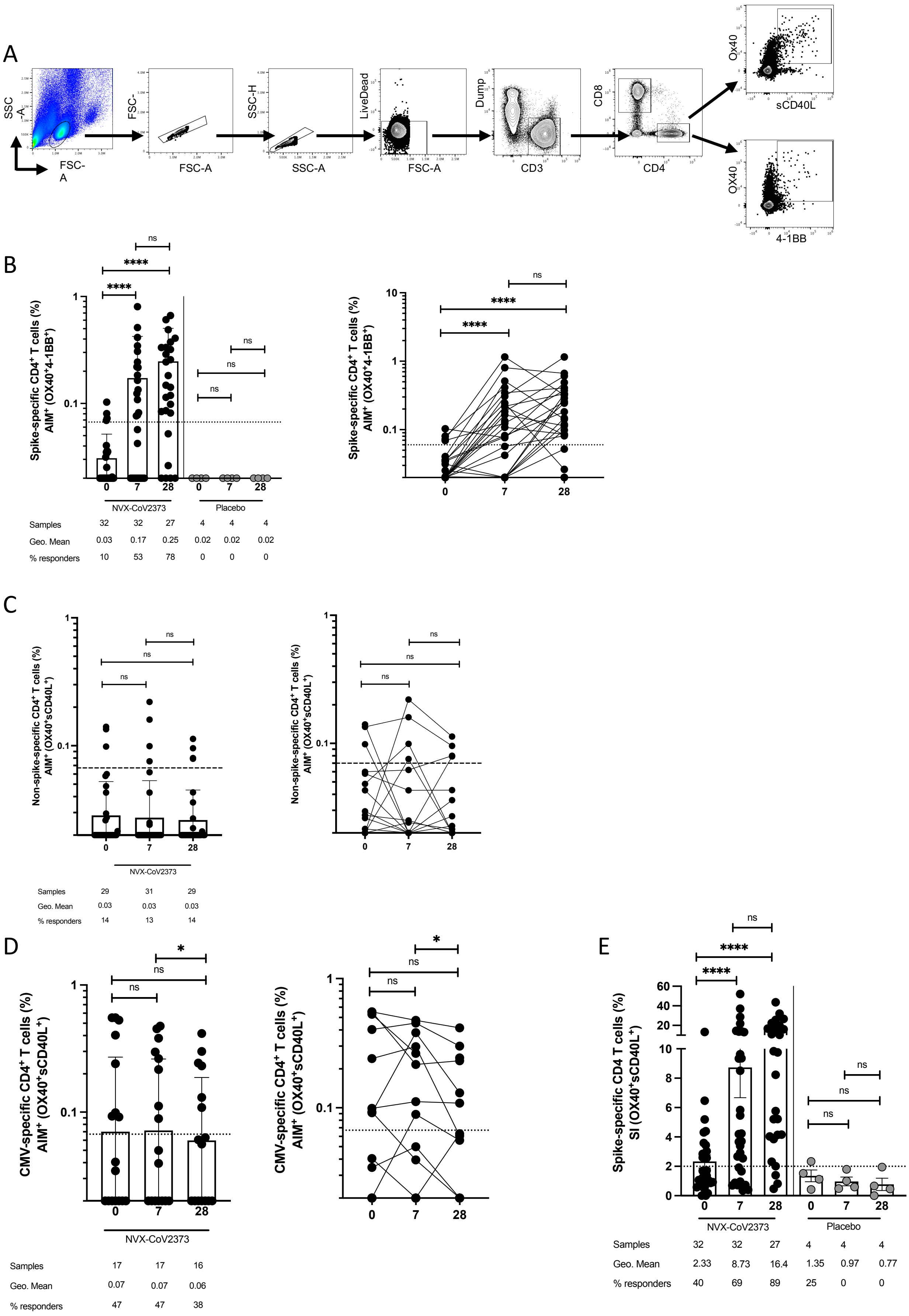

**Figure S1. (A)** FACS gating strategy used to define AIM$^+$CD4$^+$ T cells at D0, D7 and D28 post-vaccination. **(B)** Spike-specific OX40$^+$4-1BB$^+$ AIM$^+$ CD4$^+$ T cell responses in vaccinees (black dots) and placebo controls (grey dots), graphed as both grouped and paired comparisons for the vaccine cohort (**C)** SARS-CoV-2 non-spike-specific OX40$^+$CD40L$^+$ AIM$^+$ CD4$^+$ T cell responses in vaccinees at D0, 7 and 28 post-vaccination. **(D)** CMV class I/II MP-specific OX40$^+$CD40L$^+$ AIM$^+$ CD4$^+$ T cell responses in vaccinees at D0, 7 and 28 post-vaccination. **(E)** Stimulation indices for OX40$^+$CD40L$^+$ CD4$^+$ AIM$^+$ T cells in vaccinees and controls at D0, D7 and D28 post-vaccination. Dotted line indicates LOQ, and is calculated as the geometric mean of all sample DMSO wells multiplied by the geometric SD factor. % responders are calculated as responses ≥ LOQ divided by the total samples in the group.

Figure S2

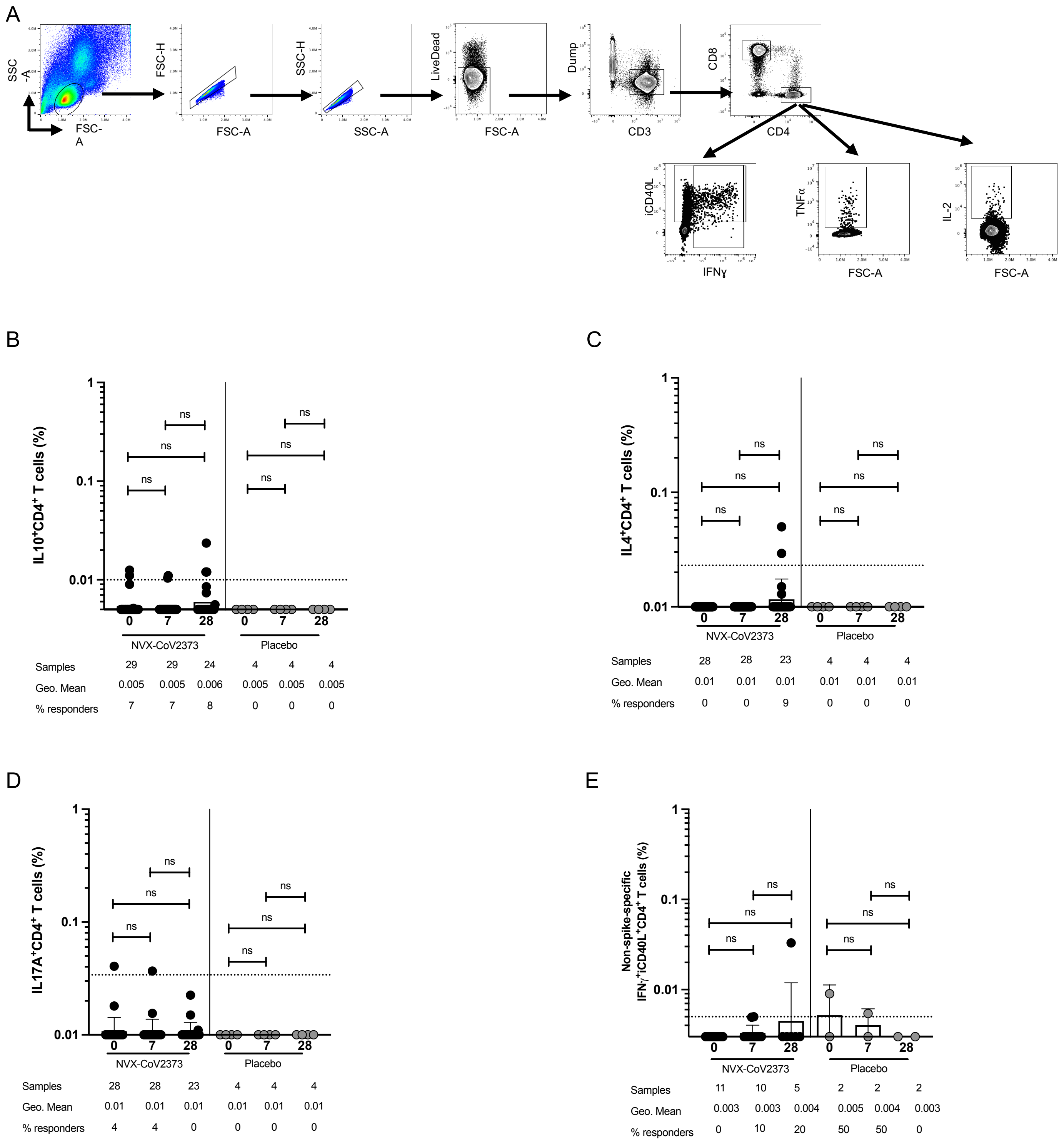
A
SSC-A
FSC-A
FSC-H
SSC-H
LiveDead
Dump
CD3
CD8
CD4
iCD40L
IFNɣ
TNFα
IL-2
B
IL10+CD4+ T cells (%)
ns
NVX-CoV2373
Placebo
Samples 29 29 24 4 4 4
Geo. Mean 0.005 0.005 0.006 0.005 0.005 0.005
% responders 7 7 8 0 0 0
C
IL4+CD4+ T cells (%)
NVX-CoV2373
Placebo
Samples 28 28 23 4 4 4
Geo. Mean 0.01 0.01 0.01 0.01 0.01 0.01
% responders 0 0 9 0 0 0
D
IL17A+CD4+ T cells (%)
NVX-CoV2373
Placebo
Samples 28 28 23 4 4 4
Geo. Mean 0.01 0.01 0.01 0.01 0.01 0.01
% responders 4 4 0 0 0 0
E
Non-spike-specific IFNγ+iCD40L+CD4+ T cells (%)
NVX-CoV2373
Placebo
Samples 11 10 5 2 2 2
Geo. Mean 0.003 0.003 0.004 0.005 0.004 0.003
% responders 0 10 20 50 50 0

**Figure S2. (A)** FACS gating strategy used to define cytokine$^+$ CD4$^+$ T cells at D0, D7 and D28 post-vaccination. Proportion of **(B)** IL10$^+$, **(C)** IL4$^+$ and **(D)** IL17A$^+$ spike-specific CD4 T cells detected following peptide stimulation. **(E)** IFNɣ$^+$CD40L$^+$ CD4$^+$ T cells measured in response to SARS-CoV-2 Remainder MP stimulation. Dotted line indicates LOQ, and is calculated as the geometric mean of all sample DMSO wells multiplied by the geometric SD factor. % responders are calculated as responses ≥ LOQ divided by the total samples in the group.

### Figure S3

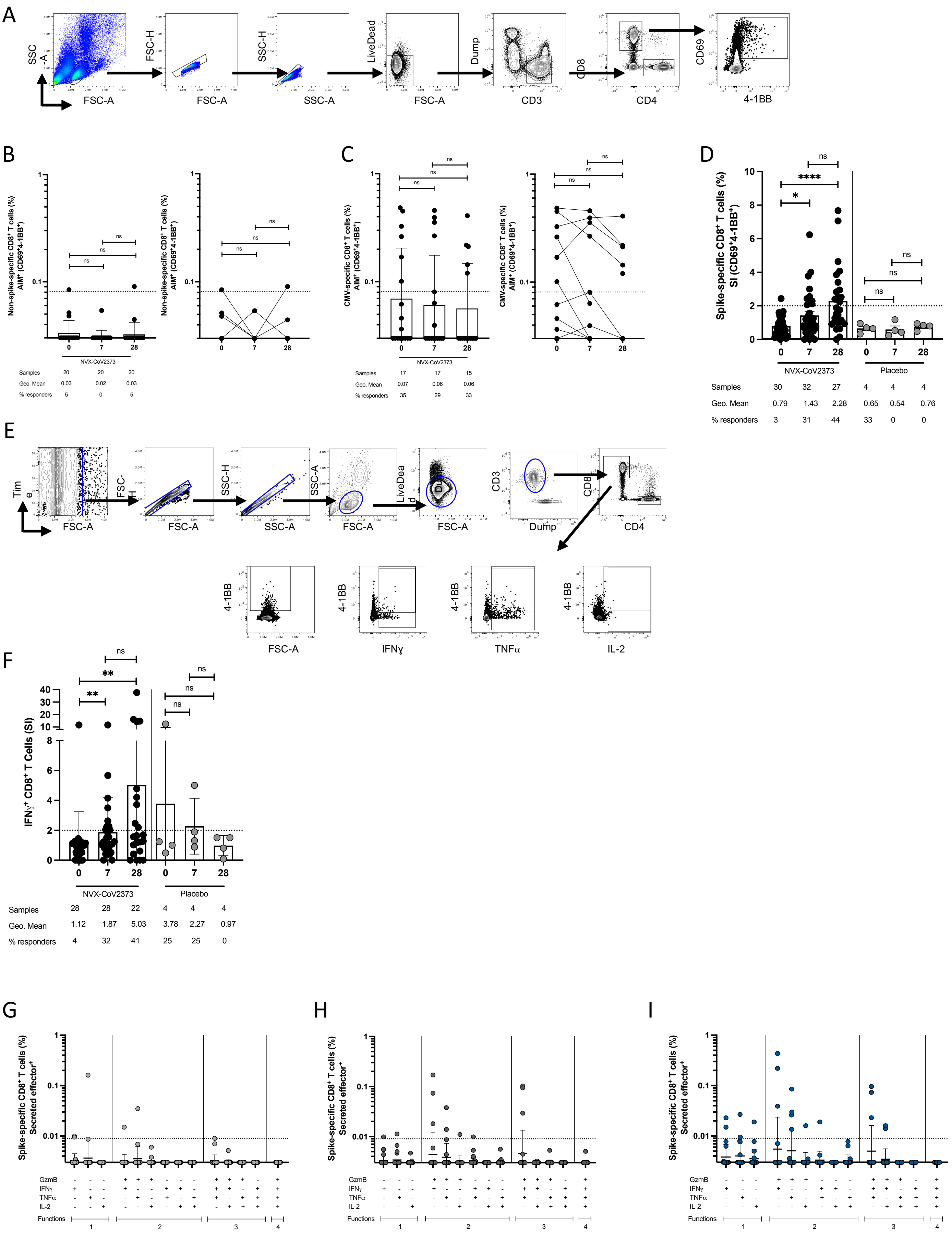
A
SSC-A
FSC-A
FSC-H
SSC-H
LiveDead
Dump
CD3
CD8
CD4
CD69
4-1BB
B
Non-spike-specific CD8⁺ T cells (%) AIM⁺ (CD69⁺4-1BB⁺)
ns
NVX-CoV2373
Samples 20 20 20
Geo. Mean 0.03 0.02 0.03
% responders 5 0 5
C
CMV-specific CD8⁺ T cells (%) AIM⁺ (CD69⁺4-1BB⁺)
Samples 17 17 15
Geo. Mean 0.07 0.06 0.06
% responders 35 29 33
D
Spike-specific CD8⁺ T cells (%) SI (CD69⁺4-1BB⁺)
NVX-CoV2373
Placebo
Samples 30 32 27 4 4 4
Geo. Mean 0.79 1.43 2.28 0.65 0.54 0.76
% responders 3 31 44 33 0 0
E
Time
Dump
CD3
CD8
CD4
4-1BB
FSC-A
IFNγ
TNFα
IL-2
F
IFNγ⁺ CD8⁺ T Cells (SI)
NVX-CoV2373
Placebo
Samples 28 28 22 4 4 4
Geo. Mean 1.12 1.87 5.03 3.78 2.27 0.97
% responders 4 32 41 25 25 0
G
H
I
Spike-specific CD8⁺ T cells (%) Secreted effector⁺
GzmB
IFNγ
TNFα
IL-2
Functions

**Figure S3. (A)** FACS gating strategy used to define AIM$^+$ CD8$^+$ T cells at D0, D7 and D28 post-vaccination. **(B)** SARS-CoV-2 Remainder MP-specific CD69$^+$4-1BB$^+$ AIM$^+$ CD8$^+$ T cell responses in vaccinees at D0, 7 and 28 post-vaccination. **(C)** CMV class I/II MP-specific CD69$^+$4-1BB$^+$ AIM$^+$ CD8$^+$ T cell responses in vaccinees at D0, 7 and 28 post-vaccination. **(D)** Stimulation indices for CD69$^+$4-1BB$^+$ CD8$^+$ AIM$^+$ T cells in vaccinees and controls at D0, D7 and D28 post-vaccination. (E) Gating strategy used to identify cytokine-producing CD8$^+$ T cells. **(F)** IFNɣ$^+$CD8$^+$ T cells measured in response to SARS-CoV-2 non-spike MP stimulation. Predominant multifunctional profiles of spike-specific CD8$^+$ T cells with one, two, three or four functions were analyzed at **(I)** D0, **(J)** D7, and **(K)** D28 post vaccination. Dotted line indicates LOQ for the assay, and is calculated as the geometric mean of all sample DMSO wells multiplied by the geometric SD factor. % responders are calculated as responses ≥ LOQ divided by the total samples in the group.

Figure S4

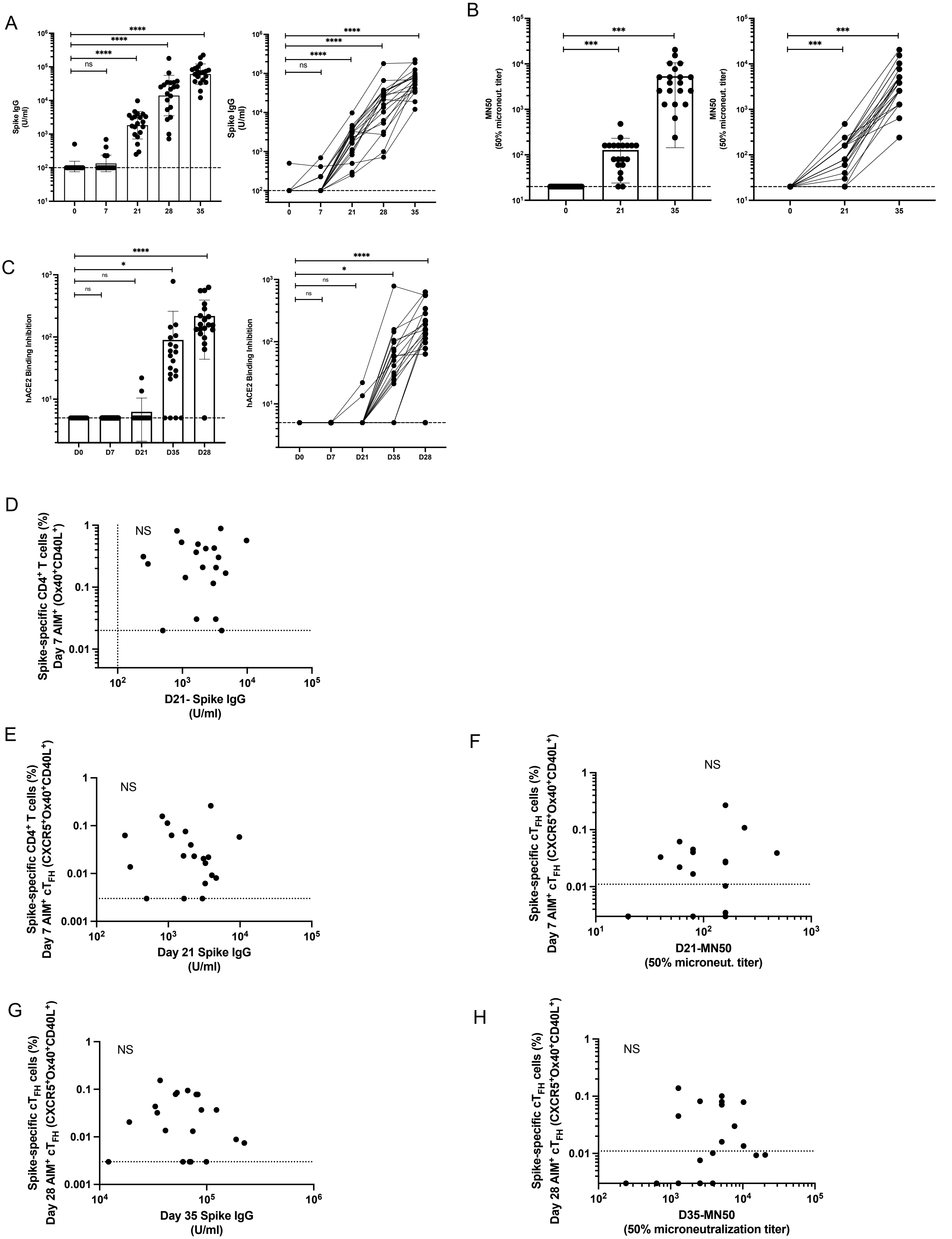

A
Spike IgG (U/ml)
0
7
21
28
35
ns
****
B
MN50 (50% microneut. titer)
***
C
hACE2 Binding Inhibition
D0
D7
D21
D35
D28
*
D
Spike-specific CD4+ T cells (%) Day 7 AIM+ (Ox40+CD40L+)
D21- Spike IgG (U/ml)
NS
E
Spike-specific CD4+ T cells (%) Day 7 AIM+ cTFH (CXCR5+Ox40+CD40L+)
Day 21 Spike IgG (U/ml)
F
Spike-specific cTFH cells (%) Day 7 AIM+ cTFH (CXCR5+Ox40+CD40L+)
D21-MN50 (50% microneut. titer)
G
Spike-specific cTFH cells (%) Day 28 AIM+ cTFH (CXCR5+Ox40+CD40L+)
Day 35 Spike IgG (U/ml)
H
D35-MN50 (50% microneutralization titer)

**Figure S4. (A)** Spike IgG titers, **(B)** SARS-CoV-2 neutralization titers, and **(C)** hACE2 binding inhibition measured in vaccinees at D0, 7, and 28 post-immunization. **(D)** Correlation of D7 SARS-CoV-2 Spike-specific $AIM^+$ $CD4^+$ T cells and D21 spike-specific IgG. Spike IgG and neutralizing antibody correlations with $AIM^+$ $cT_{FH}$ cells at **(E, F)** D7 post-1$^{st}$, or **(G, H)** D7 post-2$^{nd}$ immunization.

Figure S5

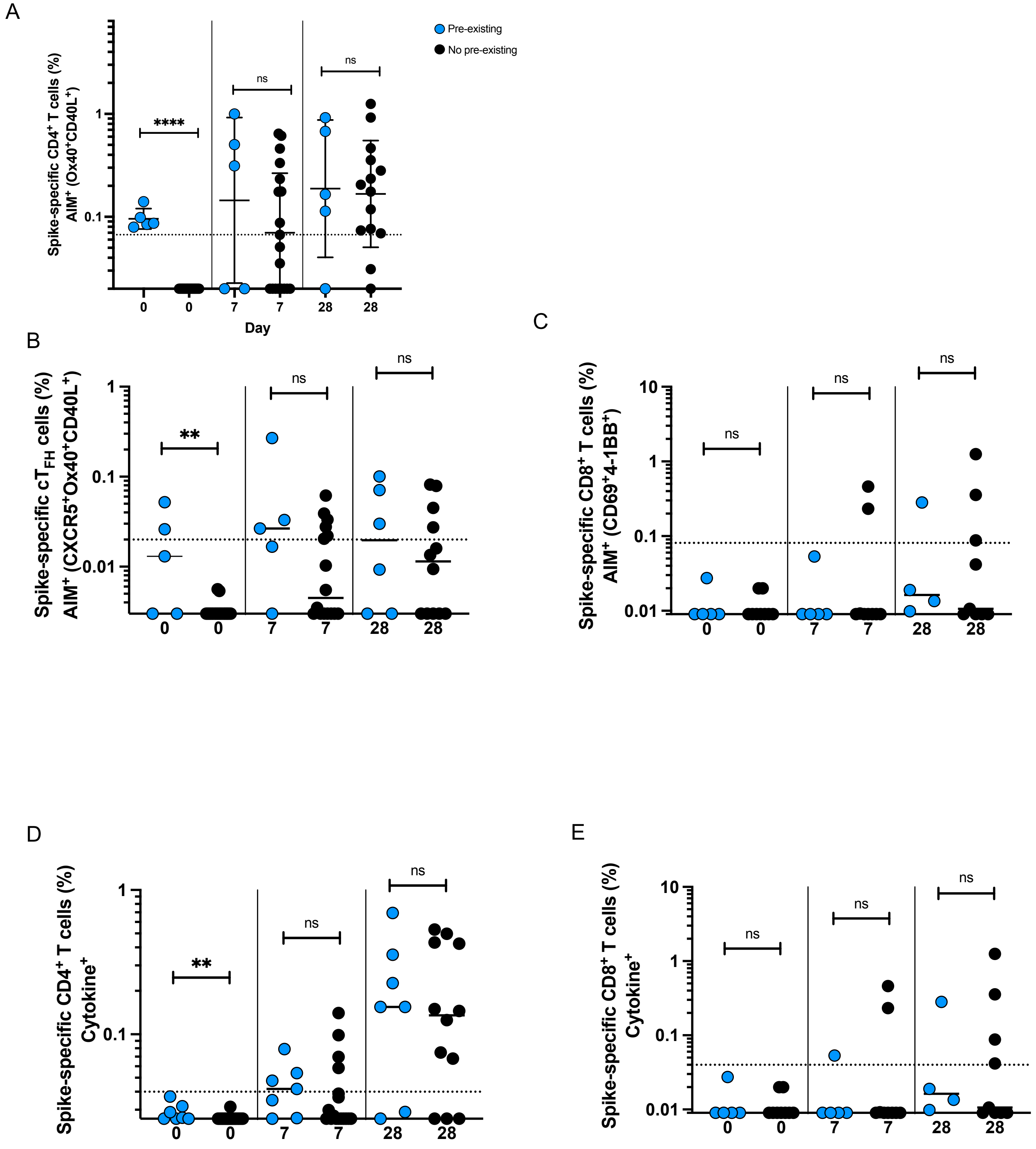
A
Pre-existing
No pre-existing
Spike-specific CD4+ T cells (%)
AIM+ (Ox40+CD40L+)
****
ns
ns
1
0.1
0
0
7
7
28
28
Day
B
Spike-specific cTFH cells (%)
AIM+ (CXCR5+Ox40+CD40L+)
**
ns
ns
1
0.1
0.01
0
0
7
7
28
28
C
Spike-specific CD8+ T cells (%)
AIM+ (CD69+4-1BB+)
ns
ns
ns
10
1
0.1
0.01
0
0
7
7
28
28
D
Spike-specific CD4+ T cells (%)
Cytokine+
**
ns
ns
1
0.1
0
0
7
7
28
28
E
Spike-specific CD8+ T cells (%)
Cytokine+
ns
ns
ns
10
1
0.1
0.01
0
0
7
7
28
28

**Figure S5. A)** Spike-specific $CD4^+$ $AIM^+$ T cells at baseline and post-vaccination grouped according to donors with $CD4^+AIM^+$ T cell responses at D0 above the LOS for the assay (blue dots) or no detectable $CD4^+$ $AIM^+$ T cell responses at D0 (black dots). Effect of high pre-existing $CD4^+$ $AIM^+$ T cells on **(B)** spike specific $AIM^+$ $cT_{FH}$ cells, **(C)** spike-specific $AIM^+$ $CD8^+$ T cells, **(D)** total Spike-specific $CD4^+$ T cell and **(E)** spike-specific $CD8^+$ T cell cytokine production. Dotted line indicates LOQ for the assay, and is calculated as the geometric mean of all sample DMSO wells multiplied by the geometric SD factor.
